# The IL-1β–BHLHE40 Axis in the Pathogenesis of Kawasaki Disease: Insights from Cell-Type-Specific Expression Analysis

**DOI:** 10.64898/2026.08.31.748192

**Authors:** Sho Ishigaki, Tatsuya Koreeda, Hiroshi Honda

**Affiliations:** Kawasaki Municipal Kawasaki Hospital, Kanagawa, Japan; CLINIC FOR Group, Tokyo, Japan; Honda Biotech Laboratory, Tochigi, Japan; University of Yamanashi, Yamanashi, Japan

**Keywords:** Kawasaki disease, BHLHE40, interleukin-1, transcriptomics, vasculitis

## Abstract

Kawasaki disease (KD) is an acute systemic vasculitis of unknown etiology that mainly affects infants and young children and can cause coronary artery dilatation or aneurysm formation. Although early treatment has improved outcomes, disease-specific biomarkers and the immune mechanisms that sustain vascular inflammation remain incompletely defined. Innate immune activation, particularly interleukin-1 (IL-1) signaling, has been implicated in KD pathogenesis, and microbial component-induced mouse models reproduce key features of KD-like coronary arteritis. BHLHE40 is an inflammation-associated transcription factor that regulates cytokines such as GM-CSF, IFN-γ, and IL-10, suggesting a potential connection between IL-1 signaling and downstream effector programs in KD.

In this study, we integrated publicly available single-cell and bulk transcriptomic datasets from human KD and LCWE-induced murine vasculitis. Human analyses included comparisons with healthy or febrile controls, whereas the mouse analyses evaluated vascular cell-type expression and the response to IL-1 receptor blockade. IL1B and BHLHE40 were increased in independent whole-blood KD cohorts, with modest but statistically significant positive correlations between their expression levels. Single-cell analysis showed that IL1B was concentrated in myeloid populations, whereas BHLHE40 was distributed across several immune populations, including NK and T cells. In LCWE-treated mice, vascular Bhlhe40 expression was reduced by Anakinra, and Il1b and Bhlhe40 were strongly correlated within the LCWE group.

These findings identify BHLHE40 as a candidate transcriptional readout of IL-1-associated inflammation in KD and support a working model in which IL-1-rich innate immune activation is linked to BHLHE40-associated effector programs in vascular immune cells.

## Introduction

Kawasaki disease (KD) is an acute systemic vasculitis of unknown etiology that predominantly affects infants and young children and can lead to coronary artery dilatation, aneurysm formation, and long-term cardiovascular morbidity [1,2]. Although early diagnosis and treatment with intravenous immunoglobulin have substantially improved clinical outcomes, KD remains clinically challenging because disease-specific biomarkers are lacking, a subset of patients is resistant to initial therapy, and the mechanisms that sustain vascular inflammation are incompletely understood [1,3]. Defining molecular programs that are shared between circulating inflammation and affected vascular tissue may therefore improve both mechanistic understanding and biomarker development.

Recent studies have suggested that KD involves dysregulated innate immune activation triggered by infectious or environmental stimuli [2,3]. Pattern-recognition pathways activated by pathogen-associated molecular patterns and damage-associated molecular patterns have been proposed to initiate and amplify KD-associated inflammation [2]. In experimental models, microbial components such as Candida albicans water-soluble fraction and Lactobacillus casei cell wall extract (LCWE) can induce KD-like coronary arteritis, supporting the concept that innate immune activation contributes to vascular inflammation in KD [4,5].

Among inflammatory cytokines implicated in KD, interleukin-1 (IL-1) has emerged as a key mediator of experimental KD-like vasculitis. In the LCWE-induced mouse model, IL-1β production depends on inflammasome-related pathways, and genetic or pharmacological inhibition of IL-1 signaling reduces coronary and aortic inflammation [6,7]. The translational relevance of this pathway is further supported by clinical investigation of the IL-1 receptor antagonist Anakinra in intravenous immunoglobulin-resistant KD [16]. These observations provide a rationale for identifying transcriptional programs that accompany IL-1-driven inflammation and that may bridge innate immune activation to downstream vascular effector responses.

BHLHE40 is a basic helix-loop-helix transcription factor that shapes inflammatory cytokine programs in multiple immune cell lineages. Previous studies have shown that BHLHE40 promotes pathogenic T-cell responses by supporting GM-CSF and IFN-γ production while restraining IL-10, and it has also been implicated in pro-inflammatory macrophage programs [8–10]. An IL-1-BHLHE40 relationship has been demonstrated in pathogenic T helper cells in experimental autoimmune neuroinflammation [11]. Because *IL1B* and *BHLHE40* may be expressed by different cellular compartments, their relationship in KD may represent an intercellular inflammatory circuit rather than simple co-expression within the same cell. However, whether BHLHE40 is associated with IL-1-related inflammation in human KD and in affected vascular tissue has not been systematically examined. We therefore reanalyzed complementary human and mouse transcriptomic datasets to define the cellular distribution of *IL1B* and *BHLHE40* and to determine whether vascular Bhlhe40 expression changes after IL-1 receptor blockade.

## Materials and Methods

### Datasets and Study Design

Human single-cell data were obtained from GSE200743 [12]. The deposited dataset available for analysis comprised peripheral-blood white blood cells from three patients with acute KD sampled before intravenous immunoglobulin treatment and two febrile controls. For independent whole-blood validation, GSE68004 [13] was analyzed using 76 complete KD samples and 37 healthy-control samples generated on the Illumina HumanHT-12 v4 expression array. GSE178491 [14], a whole-blood RNA-sequencing cohort, was analyzed using the NCBI-generated RNA-sequencing count matrix available through GEO2R. The analysis included 105 acute-phase KD samples and 29 non-KD febrile-control samples represented in this matrix. One additional febrile-control sample (GSM5392719) was not represented in the NCBI-generated count matrix and therefore could not be included. Convalescent KD samples and samples generated using poly(A) capture were excluded to maintain an acute-phase cross-sectional comparison and minimize technical heterogeneity. Murine bulk transcriptomic data were obtained from GSE141072 [7], which profiled abdominal aortas from 5-week-old male and female mice assigned to PBS, LCWE, or LCWE plus Anakinra groups, with four or five mice per sex-treatment group. The present analysis focused on Il1b and Bhlhe40 expression across the three treatment conditions and on their correlation within LCWE-treated animals.

Processed expression matrices and accompanying sample metadata were downloaded from the Gene Expression Omnibus. Analyses were performed within each dataset on the deposited expression scale; datasets were not merged because they differed in platform, specimen type, and cohort structure. Gene-level values were linked to the corresponding clinical or treatment metadata, and IL1B/BHLHE40 (human) or Il1b/Bhlhe40 (mouse) expression was compared between the prespecified groups. This within-dataset strategy was used to seek reproducibility across independent cohorts while minimizing assumptions introduced by cross-platform batch correction.

### Single-Cell RNA-seq Analysis

For human cell-type analysis, GSE200743 [12] was reanalyzed using the deposited single-cell expression matrices and sample annotations. Data had been generated on the 10x Genomics platform from peripheral-blood white blood cells with greater than 90% viability. For murine vascular cell-type analysis, GSE178765 [15] was reanalyzed. This dataset was generated with 10x Genomics 3′ v3 chemistry from abdominal aortas collected 2 weeks after injection; the PBS sample comprised a pool of nine aortas and the LCWE sample a pool of seven aortas. Quality filtering, normalization, dimensionality reduction, and clustering were performed before visualization on UMAP embeddings. *IL1B*/*BHLHE40* or *Il1b*/*Bhlhe40* expression was evaluated globally and across annotated cell populations. Because the human dataset included only five subjects and the murine conditions each comprised one pooled sample, cell-level plots were interpreted as descriptive evidence of cellular localization rather than as a substitute for subject-or animal-level biological replication.

## Statistical Analysis

Statistical analyses of bulk transcriptomic data were performed using GraphPad Prism version 10.0 (GraphPad Software, San Diego, CA, USA). For these analyses, each patient or mouse was considered an independent biological replicate. Comparisons between two independent groups in Figures 2A and 2B were performed using two-sided unpaired Student’s t tests. Comparisons among the PBS, LCWE, and LCWE plus Anakinra groups in Figure 4A were performed using one-way analysis of variance followed by Tukey’s multiple-comparison test. Pearson correlation coefficients were calculated to evaluate the relationships between IL1B and BHLHE40 expression within the KD samples in the two human cohorts and between Il1b and Bhlhe40 expression within the LCWE-treated mice, as shown in Figures 2C, 2D, and 4B. For the single-cell RNA-sequencing analyses shown in Figures 1 and 3, cells were visualized at the cell level but were not treated as independent biological replicates. Because the human single-cell dataset included only three patients with KD and two febrile controls, and because the murine single-cell dataset included only one pooled sample per condition, no between-group inferential statistical testing was performed using individual cells. These analyses were interpreted descriptively to assess the cellular distribution and cell-type localization of gene expression.

**Figure 1.**
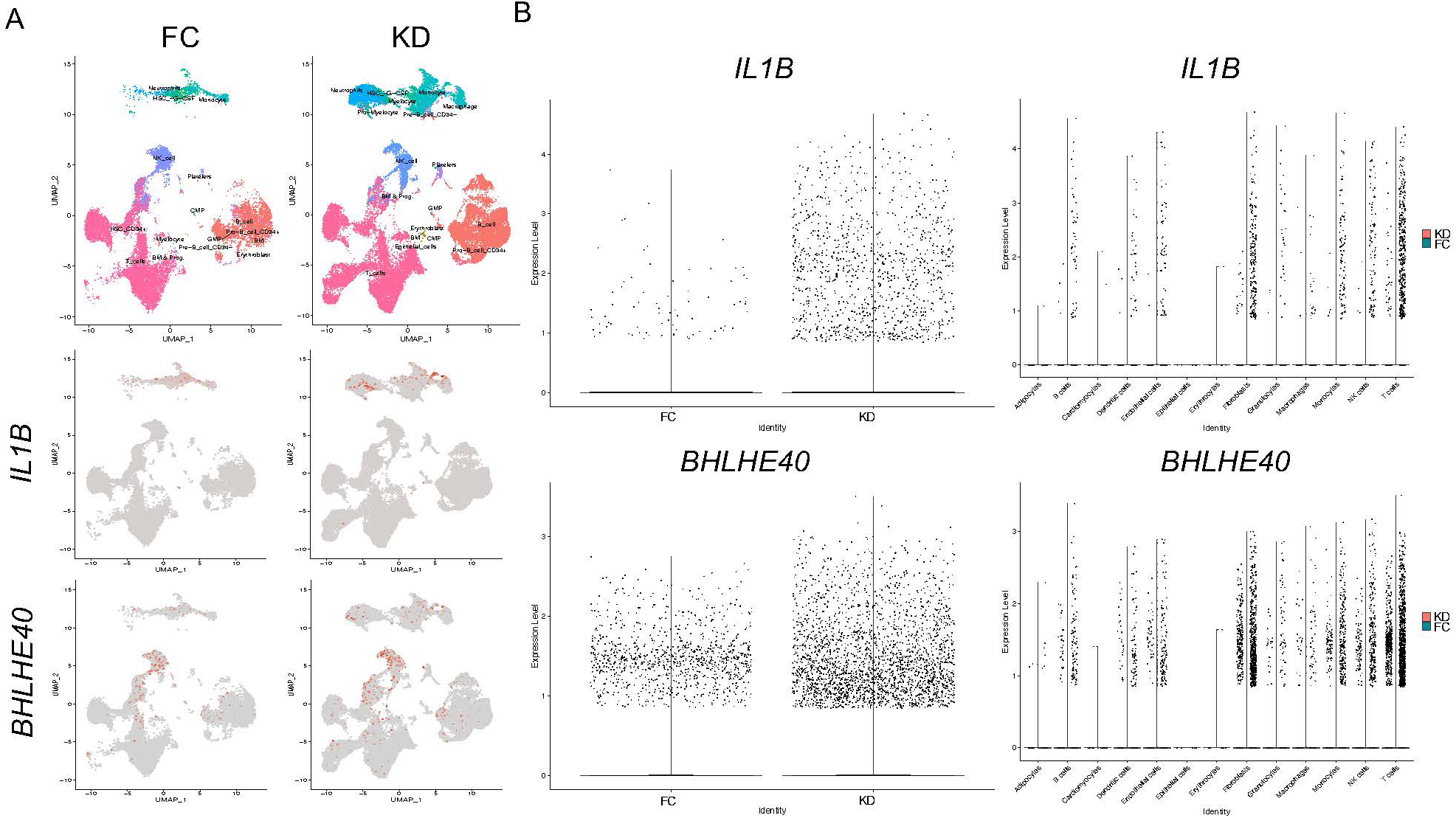
IL1B and BHLHE40 are increased in peripheral blood immune cells from patients with Kawasaki disease. Single-cell RNA-sequencing data from GSE200743 [12] were reanalyzed to evaluate *IL1B* and *BHLHE40* expression in acute KD. (A) Top, UMAP plots showing annotated peripheral-blood leukocyte populations in febrile controls and patients with KD. Bottom, feature plots showing *IL1B* and *BHLHE40* expression projected onto the corresponding UMAP embeddings. (B) Left, overall cell-level expression distributions in febrile controls and KD. Right, expression distributions stratified by annotated leukocyte population and disease group. *IL1B* was prominent in myeloid populations, whereas *BHLHE40* was distributed across several leukocyte subsets. Cell-level distributions are presented descriptively because cells were nested within subjects and were not treated as independent biological replicates. No inferential statistical testing was performed. FC, febrile control; KD, Kawasaki disease; UMAP, uniform manifold approximation and projection.

**Figure 2.**
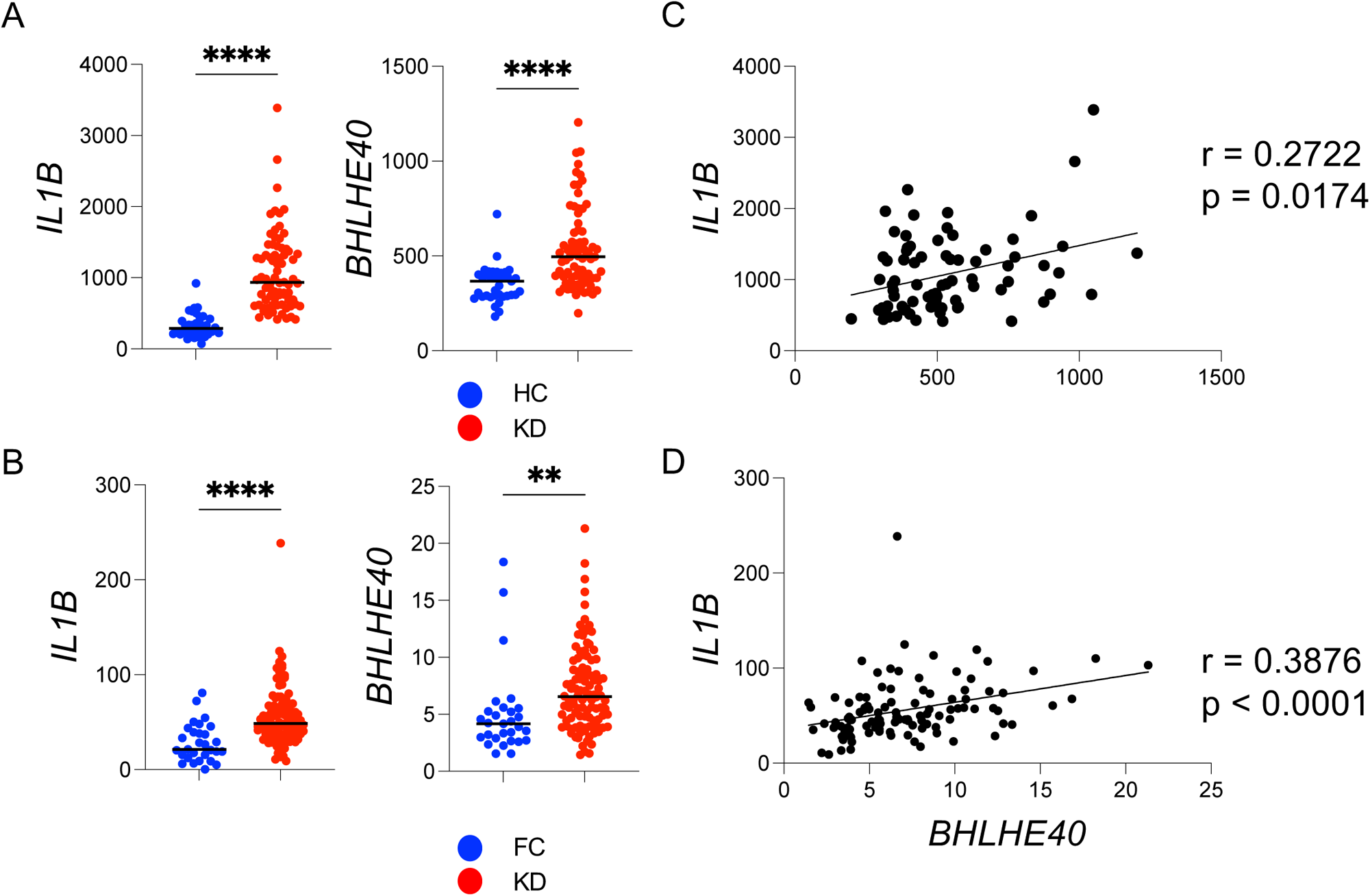
Independent human transcriptomic cohorts validate increased IL1B and BHLHE40 expression in Kawasaki disease. *IL1B* and *BHLHE40* expression was analyzed in two independent whole-blood KD cohorts. (A) Expression in healthy controls and complete KD in GSE68004 [13]. (B) Expression in non-KD febrile controls and Kawasaki disease in GSE178491 [14]. (C) Correlation between IL1B and BHLHE40 among KD samples in GSE68004. (D) Correlation between IL1B and BHLHE40 among KD samples in GSE178491. Correlation between *IL1B* and *BHLHE40* in GSE178491. Each dot represents one sample; lines indicate linear regression. Correlation coefficients and p values are shown in the plots. FC, febrile control; HC, healthy control; KD, Kawasaki disease. Group comparisons were performed using two-sided unpaired Student’s t tests. **p < 0.01; ****p < 0.0001.

**Figure 3.**
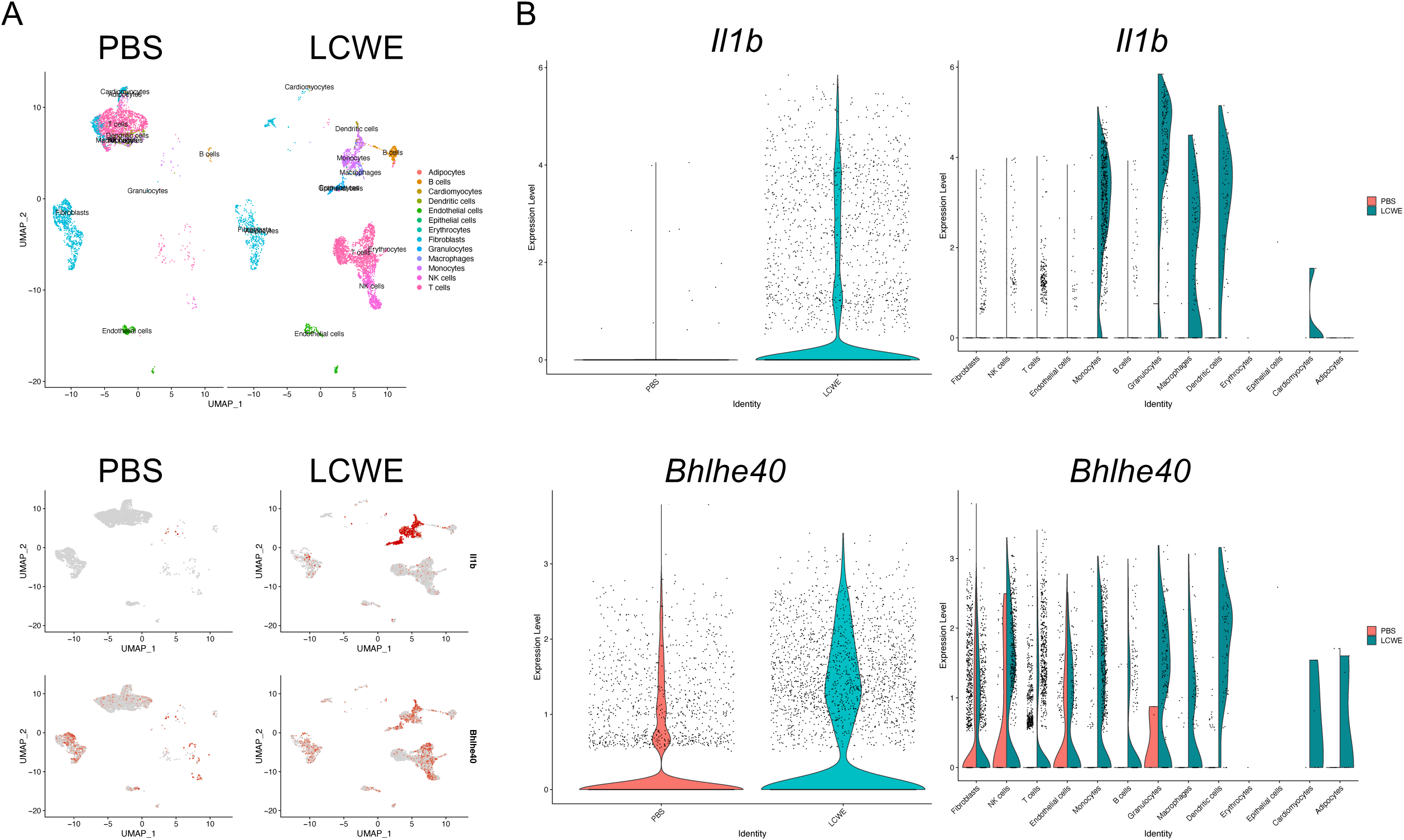
LCWE-induced Kawasaki disease vasculitis is associated with induction of *Il1b* and *Bhlhe40* in vascular immune cells. Single-cell RNA-sequencing data from abdominal aortas of PBS-or LCWE-injected mice (GSE178765) [15] were reanalyzed. The PBS sample comprised a pool of nine aortas and the LCWE sample a pool of seven aortas. (A) Top, UMAP plots showing annotated abdominal-aortic cell populations. Bottom, feature plots showing *Il1b* and *Bhlhe40* expression on the corresponding UMAP embeddings. (B) Left, overall cell-level expression distributions. Right, expression distributions stratified by annotated vascular cell population and treatment condition. *Il1b* was concentrated in inflammatory myeloid populations, whereas *Bhlhe40* was distributed across lymphoid, myeloid, and stromal subsets. No inferential statistical testing was performed because each condition was represented by a single pooled sample. LCWE, Lactobacillus casei cell wall extract; PBS, phosphate-buffered saline.

**Figure 4.**
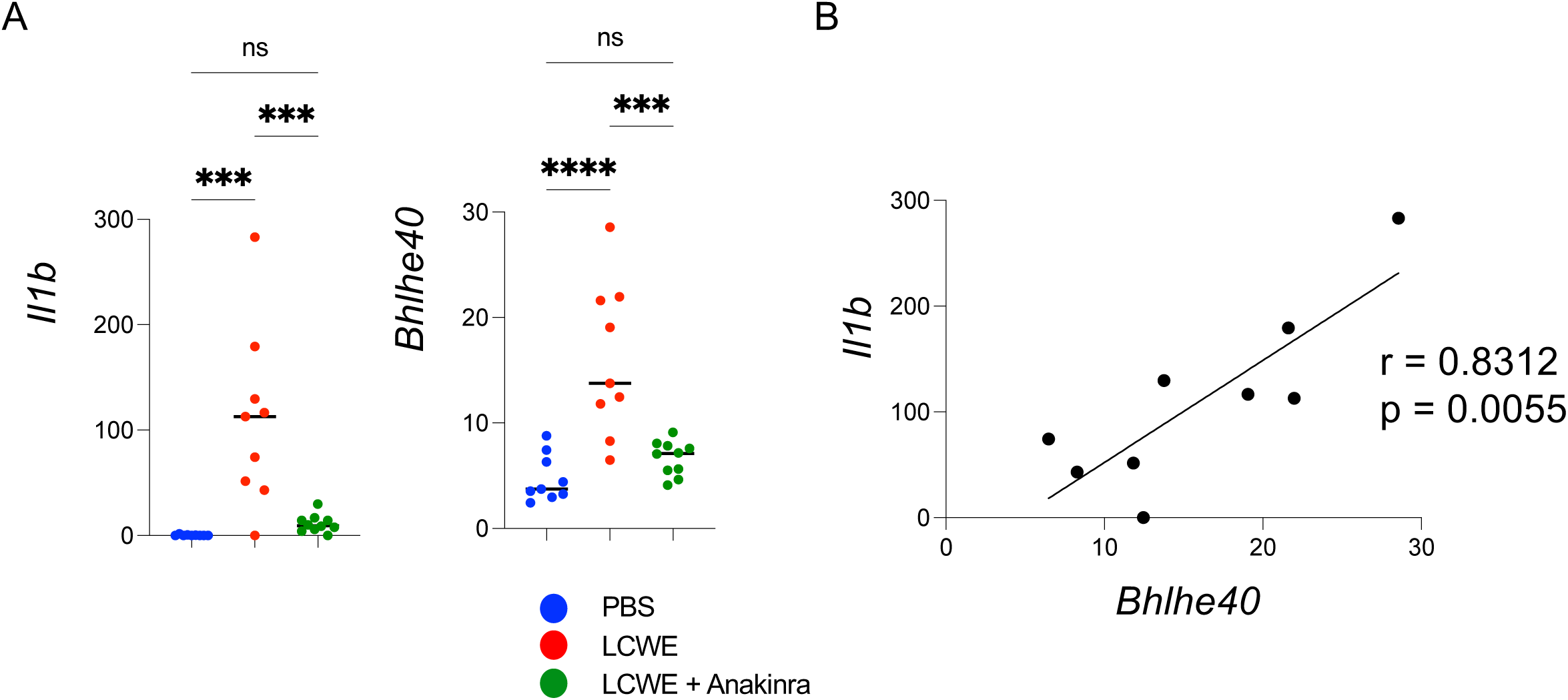
IL-1 receptor blockade suppresses LCWE-induced Bhlhe40 expression in the Kawasaki disease vasculitis model. Bulk abdominal-aorta transcriptomic data from GSE141072 [7] were reanalyzed to evaluate IL-1 receptor blockade. (A) Il1b and Bhlhe40 expression in PBS control, LCWE, and LCWE plus Anakinra groups. LCWE increased both transcripts, whereas Anakinra reduced expression toward control levels. (B) Correlation between Il1b and Bhlhe40 within LCWE-treated mice. Each dot represents one mouse; the line indicates linear regression. The correlation coefficient and p value are shown in the plot. LCWE, Lactobacillus casei cell wall extract. Group comparisons were performed using one-way analysis of variance followed by Tukey’s multiple-comparison test. ns, not significant; ***p < 0.001; ****p < 0.0001.

All statistical tests were two-sided, and p < 0.05 was considered statistically significant.

## Results

### 1. Elevated IL-1β and BHLHE40 Gene Expression in Peripheral Blood of Kawasaki Disease Patients

We first examined IL1B and BHLHE40 expression in peripheral-blood white blood cells from acute KD patients (n = 3) and FC (n = 2) in GSE200743 [12]. UMAP visualization suggested a greater representation of neutrophils in the KD samples, consistent with the prominent innate immune response of acute KD (Figure 1A). Cell-level visualization also suggested higher *IL1B* and *BHLHE40* expression in KD than in febrile controls (Figure 1B). *IL1B* signal was most prominent in myeloid populations, whereas *BHLHE40* signal was detectable across multiple leukocyte subsets. Because the dataset included only three patients with KD and two FC, these observations were interpreted descriptively and used to localize the candidate axis before evaluation in larger independent whole-blood cohorts.

### 2. Comparison of IL-1β and BHLHE40 Gene Expression in KD Versus Control Cohorts

We next assessed the reproducibility of these findings using two larger whole-blood transcriptomic datasets with different control populations. In GSE68004, *IL1B* and *BHLHE40* expression levels were significantly higher in patients with complete KD (n = 76) than in healthy controls (n = 37; Figure 2A). Similarly, in GSE178491, both genes were significantly upregulated in patients with acute-phase KD (n = 105) compared with non-KD febrile controls (n = 29; Figure 2B). These consistent findings across distinct control populations indicate that the increased expression of *IL1B* and *BHLHE40* in KD was not limited to comparisons with healthy children.

Pearson correlation analyses performed exclusively within the KD groups further demonstrated positive associations between *IL1B* and *BHLHE40* expression in both independent cohorts. A modest positive correlation was observed among patients with complete KD in GSE68004 (r = 0.2722, p = 0.0174; Figure 2C), whereas a somewhat stronger positive correlation was detected among patients with acute-phase KD in GSE178491 (r = 0.3876, p < 0.0001; Figure 2D). Because these analyses were restricted to patients with KD, the observed correlations cannot be explained solely by the overall expression differences between KD and control groups. Collectively, these findings demonstrate a reproducible, albeit modest, association between *IL1B* and *BHLHE40* expression within KD and support their coordinated involvement in the inflammatory response associated with the disease.

### 3. Single-Cell RNA Sequencing Analysis Reveals Cell-Type Specific Expression Patterns

To define the vascular cellular context of *Il1b* and *Bhlhe40*, we analyzed abdominal-aorta scRNA-seq data from PBS- and LCWE-injected mice in GSE178765 [15] (Figure 3). In the pooled LCWE sample, inflammatory immune populations were more prominently represented than in the pooled PBS sample. *Il1b* expression was concentrated in monocytes, macrophages, granulocytes, and dendritic cells. *Bhlhe40* was more broadly distributed and was readily detected in lymphoid populations, including NK and T cells, as well as in subsets of myeloid and stromal cells (Figure 3A,B). This partial separation of the principal Il1b- and Bhlhe40-expressing compartments is compatible with an intercellular relay from IL-1-producing myeloid cells to BHLHE40-expressing effector populations. Because each condition was represented by one pooled sample, these observations were interpreted descriptively and support cell-type localization, but not animal-level changes in cell abundance or gene expression.

### 4. IL-1 Inhibition by Anakinra Suppresses Bhlhe40 Expression in the KD Mouse Model

To determine whether vascular Bhlhe40 expression was responsive to interruption of IL-1 signaling, we analyzed abdominal-aorta transcriptomes from PBS-, LCWE-, and LCWE plus Anakinra-treated mice in GSE141072 [7]. LCWE increased both Il1b and Bhlhe40, whereas Anakinra reduced both transcripts toward control levels (Figure 4A). The Anakinra-treated group was not significantly different from PBS for either gene in the displayed comparisons. Within LCWE-treated mice, Il1b and Bhlhe40 were strongly correlated (r = 0.8312, p = 0.0055; Figure 4B), a substantially stronger relationship than that observed in the heterogeneous human whole-blood cohorts.

Together, these data show that Bhlhe40 expression tracks with the intensity of IL-1-associated vascular inflammation and is attenuated during pharmacological IL-1 receptor blockade. The intervention supports pathway responsiveness but does not by itself establish direct transcriptional regulation of Bhlhe40 by IL-1 in a specific cell type.

## Discussion

In this study, we identified BHLHE40 as a candidate transcription factor associated with IL-1-related inflammation in KD by integrating four complementary transcriptomic settings: human peripheral-blood single-cell data, two independent human whole-blood cohorts, murine vascular single-cell data, and a murine pharmacological intervention dataset. Across these settings, BHLHE40 was increased during KD or KD-like inflammation, occupied a cellular distribution distinct from but complementary to IL1B, and decreased after IL-1 receptor blockade. The cross-species and cross-platform concordance is the principal strength of the analysis and supports BHLHE40 as a reproducible component of an IL-1-associated inflammatory program.

The human data suggest that the association is not explained solely by comparison of acutely inflamed children with healthy controls. BHLHE40 was elevated not only in GSE68004, which used healthy controls, but also in GSE178491, which used children with non-KD febrile illnesses. This is important because many inflammatory transcripts rise nonspecifically during fever. Nevertheless, the correlations between IL1B and BHLHE40 were modest, and the present analyses do not establish diagnostic specificity. Differences in leukocyte composition, disease timing, treatment status, and upstream cytokines other than IL-1 are all likely to contribute to BHLHE40 expression. BHLHE40 should therefore be viewed as a candidate component of a multigene inflammatory signature rather than a stand-alone surrogate for IL1B.

Single-cell analysis provided a mechanistic framework for interpreting the bulk-tissue association. IL1B was concentrated in inflammatory myeloid populations, whereas BHLHE40 was detected across lymphoid and other vascular cell populations. This pattern raises the possibility that IL-1-producing neutrophils, monocytes, or macrophages condition neighboring T and NK cells to engage BHLHE40-associated effector programs. Such a model is biologically plausible because BHLHE40 promotes GM-CSF and IFN-γ while restraining IL-10 in activated T cells [8,9,11], and it can also reinforce pro-inflammatory macrophage function [10]. At the same time, ligand-receptor signaling was not directly tested here; altered cell abundance or parallel induction by other inflammatory cues could generate a similar transcriptomic pattern.

The LCWE model added a vascular-tissue dimension that is difficult to obtain from children with KD. Il1b and Bhlhe40 were both induced in inflamed aortas, and the distribution of the two genes across vascular immune populations broadly paralleled the human peripheral-blood findings. This convergence supports the utility of the LCWE model for testing the proposed circuit. However, the mouse scRNA-seq comparison was generated from one pooled specimen per condition (nine PBS aortas and seven LCWE aortas). It therefore identifies candidate cellular sources but cannot quantify between-animal variability or prove that the same cells change expression after LCWE exposure. The reduction in Bhlhe40 after Anakinra provides the strongest interventional support for the proposed relationship. This result is consistent with prior evidence that IL-1 induces Bhlhe40 in pathogenic helper T cells [11] and is clinically relevant because Anakinra has been evaluated in intravenous immunoglobulin-resistant KD [16]. Nevertheless, Anakinra can reduce the overall inflammatory burden and change vascular leukocyte composition, either of which could secondarily lower bulk Bhlhe40 expression. Demonstrating a direct pathway will require cell-type-resolved IL-1β stimulation, measurement of BHLHE40 induction kinetics, and interruption of IL-1R1 signaling in purified candidate responder populations.

These findings have potential biomarker and mechanistic implications. As a biomarker, BHLHE40 should next be tested in longitudinal samples obtained before and after intravenous immunoglobulin, in febrile disease controls, and in patients stratified by treatment response and coronary artery outcome. Its value should be assessed as an addition to, rather than a replacement for, established clinical and inflammatory variables. Mechanistically, BHLHE40 may help sustain vascular inflammation by converting an upstream innate cytokine signal into durable effector-cell programs. This possibility is especially relevant if BHLHE40 remains active after the initiating IL-1 signal has begun to decline.

A decisive functional test would combine lineage-specific perturbation with vascular phenotyping. Conditional deletion or knockdown of Bhlhe40 in T cells, NK cells, and myeloid cells could identify the compartment required for LCWE-induced inflammation. Ex vivo IL-1β stimulation of sorted vascular or peripheral immune subsets should be paired with BHLHE40 protein measurement and downstream readouts such as GM-CSF, IFN-γ, IL-10, and TNF. Rescue experiments that restore BHLHE40 in the relevant deficient lineage would further distinguish direct transcriptional control from secondary changes in cell recruitment. Linking these perturbations to coronary/aortic histology, immune-cell infiltration, and vascular cytokine production would establish whether the IL-1-BHLHE40 relationship is causal rather than merely associative.

Several limitations should be acknowledged. First, this study was a retrospective reanalysis of publicly available datasets and was therefore constrained by deposited metadata, heterogeneous platforms, differences in sample processing, and nonuniform clinical covariates. Second, the human single-cell dataset contained only three KD patients and two febrile controls, and the murine single-cell dataset contained one pooled sample per condition; cell-level observations must not be interpreted as independent patient-or animal-level replication. Third, the Anakinra dataset included both sexes, but the present focused analysis did not formally model sex-by-treatment interactions. Fourth, bulk expression cannot distinguish cell-intrinsic transcriptional regulation from shifts in cellular composition. Finally, the study is correlative at the cellular level and does not establish that BHLHE40 is required for coronary or aortic inflammation.

In conclusion, integrated human and mouse transcriptomic evidence links BHLHE40 to IL-1-associated inflammation in KD. Its reproducible elevation in human KD, localization to defined vascular immune compartments, suppression after IL-1 receptor blockade, and correlation with IL1B support BHLHE40 as a candidate transcriptional readout and potential effector of KD-associated vascular inflammation. Cell-type-specific perturbation studies and longitudinal patient validation are now required to distinguish biomarker association from causal function.

